# Biomineralization from platelet δ-granules as the origin of cardiovascular calcification in humans and other animals

**DOI:** 10.64898/2026.08.01.742085

**Authors:** Sergio Bertazzo, Elena Tsolaki, Shweta Agarwal, Najma Latif, Ann McCormack, Padmini Sarathchandra, Magdi H Yacoub, Inge K Hermann, Ken Smith, Janice Tsui, Adrian H Chester

**Affiliations:** Department of Medical Physics & Biomedical Engineering, University College London, London WC1E 6BT, UK; London Centre for Nanotechnology, University College London, London WC1E 6BT, UK; Magdi Yacoub Institute, Imperial College London, Heart Science Centre, Harefield Hospital, Harefield, Middlesex UB9 6JH, UK; Department Materials Meet Life, Swiss Federal Laboratories for Materials Science and Technology (Empa), Lerchenfeldstrasse 5, CH-9014, St. Gallen, Switzerland; Nanoparticle Systems Engineering Laboratory, Department of Mechanical and Process Engineering, ETH Zurich, Sonneggstrasse 3, 8092 Zurich, Switzerland; Pathology and Pathogen Biology Pathobiology and Population Sciences, The Royal Veterinary College, Hawkshead Lane, North Mymms, Hertfordshire AL9 7TA UK; Department of Surgery, University College London, London WC1E 6BT, UK

## Abstract

Cardiovascular calcification is present in practically all cardiac diseases, which are the top killers in the world today^1^, and is particularly associated with atherosclerosis^2^, aortic stenosis^3^ and rheumatic fever^4^. If not the direct cause of death, calcification contributes considerably to complications that can lead to heart failure^5^. Nonetheless, the origins and mechanisms of cardiovascular calcification are still strongly debated^3,6–12^. Just over a decade ago, it has been reported that nano and micron-sized calcified spherical particles, formed from a single crystal of magnesium-containing calcium phosphate, were the first calcified structure that could be detected in cardiovascular tissue^13^. These particles were found even before any sign of cardiac disease was present and were present in all stages of cardiac diseases^13^. The ubiquity of these particles suggests their importance for the origins and development of cardiovascular calcific diseases. Here, we show that these particles originate from platelet δ-granules and are present in mammals, birds and lizards. Based on our results, we suggest a new mechanism for the origins of these particles, complementing existing models of cardiovascular calcification^7,14^, and bringing a new, early, and hitherto unaccounted key event in the process of cardiovascular calcification. This new mechanism model, along with a better understanding of the early stages of cardiovascular calcification, could open the path for the development of pharmacological prevention and treatment solutions for several cardiac diseases.

---

The origins of cardiovascular calcification in association with cardiovascular diseases^5^ have been debated for decades, and generated several proposed origin and formation mechanisms models^3,5–12,14,15^. Just over a decade ago, electron microscopy analysis revealed that cardiovascular calcification is formed from three distinct structures: highly crystalline calcified spherical particles; calcified fibres; and what was then called compact calcification^13^. The presence of these three distinct structures suggests that more than one calcification mechanism may be responsible for the origins and/or development of cardiovascular calcification. From these three structures, calcified particles were the first calcified structure that could and still can be detected in any cardiovascular tissue, well before any sign of any disease of the cardiovascular system^13^. This finding suggests that the identification of the origins and mechanism of formation of calcified particles could shed light on early and key processes that would lead to pathological cardiovascular calcification.

Whilst calcified fibres and compact calcification have several characteristics similar to calcification produced in *in vitro* and *in vivo* models^14–16^, to the best of our knowledge no model has hitherto been able to explain or even reproduce any of the unique characteristics of the calcified particles, including their morphology, size, composition and crystallinity. To identify the origins of these calcified particles, we initially determined where in the tissue most of these particles are present.

Histological slides of aortic tissue from organ donors’ samples deemed suitable for aortic valve transplant, without any sign of calcification or any other lesion (Supplementary info Table S1 for patient information) were stained with Hematoxylin and Eosin (H&E) and imaged by optical microscopy and scanning electron microscopy (SEM). Although not visible on the optical microscopy images (Fig. 1a 1bI), SEM images (Fig. 1bII and c) clearly show that the calcified particles are present and distributed all over the tissue samples (Fig. 1bII and c). No calcification, other than the calcified particles, was present in the tissue, in line with the expectation that these structures would be the first to be formed in the vascular tissue^13^. Elemental analysis (by Energy-dispersive X-ray spectroscopy (EDS)) shows that these particles are composed of calcium phosphate, with small amounts of magnesium (Fig. 1d). Transmission electron microscopy and selective area electron diffraction (TEM-SAED) (Fig. 1e), confirm that the particles observed are the same calcified particles (diffracting as a single crystal of whitlockite) described previously in different cardiovascular tissues associated to atherosclerosis, aortic stenosis and rheumatic fever^13,14,17^.

**Figure 1:**
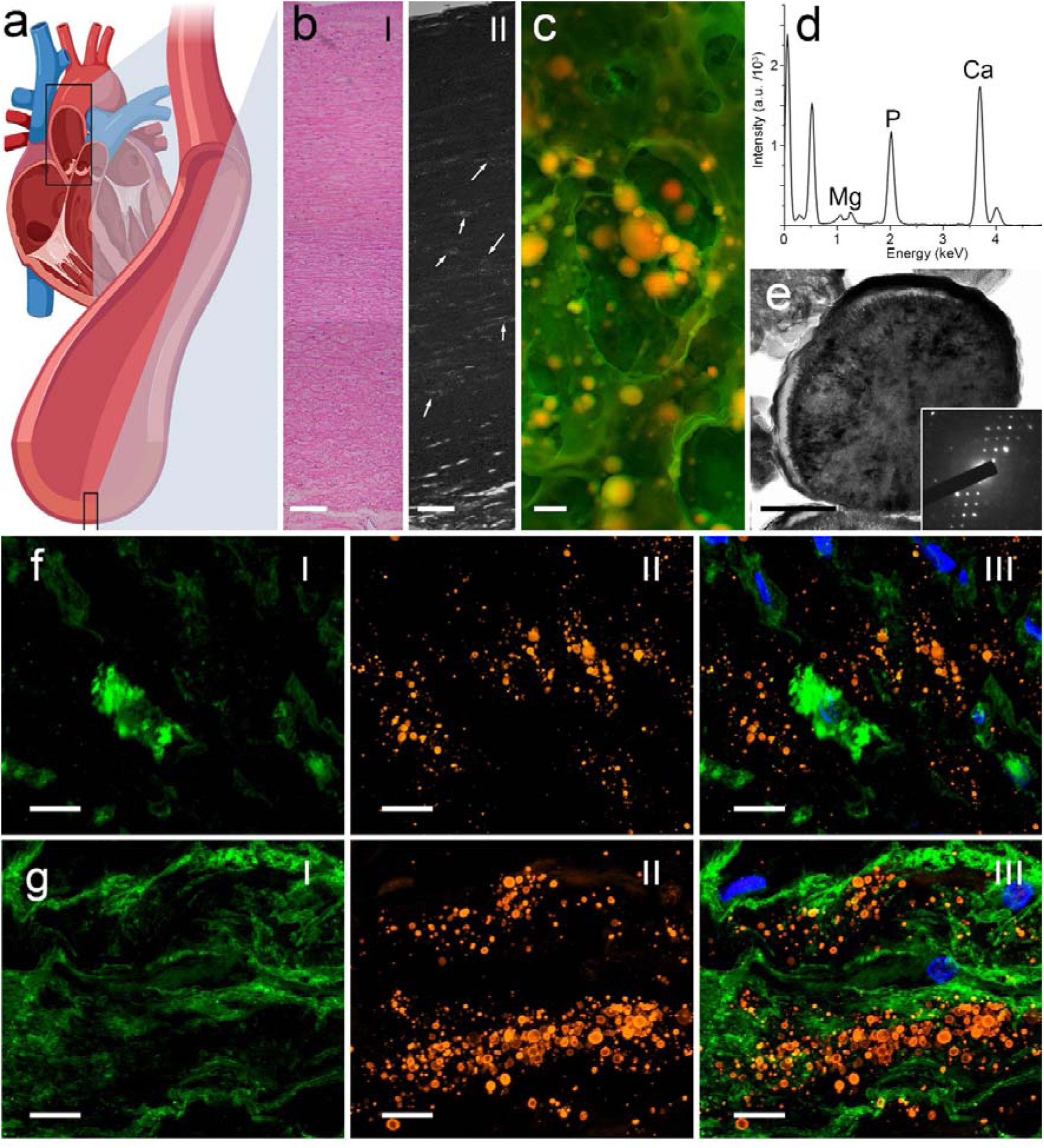
Vascular nano and micro calcified particles in aorta. **a** Diagram of sectioned aorta to be imaged by optical and electron microscopy. **b** Optical micrograph of aorta wall stained with Haematoxylin and Eosin (**I**) and SEM of the aortic wall (**II**), showing the distribution of calcified particles (some of the particles are indicated by arrows). Scale bar = 100 µm. **c Density Dependent Colour - Scanning Electron Microscope (DDC-SEM)** high magnification image of the calcified particles present in arterial tissue. Scale bar = 1 µm. **d** EDS spectrum example of calcified particles presented in **c**. **e** TEM micrograph and SAED of calcified particle sectioned by Focused Ion Beam (FIB), including the internal structure and the typical electron diffraction pattern (insert) of a single crystal of whitlockite. Scale bar = 0.2 µm. **f** Fluorescence micrograph of vascular smooth cell actin (**I**), calcified particles stained by OsteoSense (OsteoSense 680EX) (**II**) and overlay micrographs with nuclei in blue (**III**). Scale bar = 10 µm. **g** Fluorescence micrograph of collagen (**I**), calcified particles stained by OsteoSense (OsteoSense 680EX) (**II**) and overlay micrographs with nuclei in blue (**III**). Scale bar = 10 µm.

Following confirmation of the presence of the particles in the whole aortic wall, we set out to evaluate a possible connection among these calcified particles, vascular smooth muscle cells and the main extracellular matrix (ECM) component in the vasculature (collagen). Histological slides of aortic tissue were fluorescently-tagged for visualization of vascular smooth muscle alpha actin (Fig. 1f), collagen (Fig. 1g) and calcification (Fig. 1fII and 1gII). The fluorescent images show no significant colocalization of the calcified particles with either vascular smooth muscle cells (Fig. 1fIII) or collagen (Fig. 1gIII). These results firstly indicate that not all calcification stems from vascular smooth muscle cells, unlike suggested by several of the models available in literature^7,14–16^. Furthermore, the fact that the calcified particles are not associated with collagen is an indication that these particles do not derive from an osteogenic (where collagen would be strongly associated to calcification) process, which has also been suggested^5,18^.

From the insights obtained into the association among the calcified particles, vascular cells and the extracellular matrix (ECM), even if combined with the localization of the calcified particles in the tissue, we cannot yet infer the origins of these particles. Therefore, looking at the morphology, size, elemental composition, localization of the calcified particles, as well as their lack of association with vascular cells and collagen, we have been prompted to look for other cellular or extracellular components that could possibly act as templates for the formation/precipitation of these particles.

It came to our attention that platelets have been strongly associated to the onset and progression of cardiovascular diseases, and that they have even been directly associated to cardiovascular calcification^19,20^. Moreover, increased amounts of circulating platelet microvesicles have also been correlated to cardiovascular calcification^20^. Not only platelet microvesicles and exosomes released by platelets (also called platelet particles, extracellular vesicles or platelet granules), but also whole platelets^21,22^, continually pass through blood vessel walls^23,24^. These can be found in the synovial fluid^24–26^ and in several different tissues^24,27,28^. Finally, platelet vesicles are stable, being able to survive intact outside platelets^29,30^ for a considerable amount of time^31,32^.

TEM images of unstained whole platelets clearly show one type of these microvesicles, the δ-granules (also known as dense granules), which appear as electron dense spherical particles (Fig. 2a and 2b). Their high electron density is caused by large concentrations of calcium^33–35^, phosphorus^33,34,36,37^ (in the form of ADP, ATP, pyrophosphate, and polyphosphate) and magnesium^38–40^ (Fig. 2c). The diameter of δ-granules measured inside platelets (as recorded in TEM micrographs) ranges from 41 ± 2.3 nm to 436 ± 2.3 nm (Fig. 2d). Interestingly, the smallest size of δ-granules measured is virtually identical to the smaller calcified particles (38 ± 2.3 nm), as measured directly in histological slides of vascular tissue samples (Fig. 2d). Taking together the information obtained about elemental composition, morphology, and size of δ-granules, and based on two facts: first, that δ-granules (or even platelets) are present inside the vasculature^21–23^ (within the ECM); and second, that the calcified spherical particles are the first structure that can be identified in the vascular tissue (an indication that the particles might have a non-pathological origin), we hypothesize that the calcified particles may actually be δ-granules that have calcified over time, after being trapped in the ECM of the vasculature.

**Figure 2:**
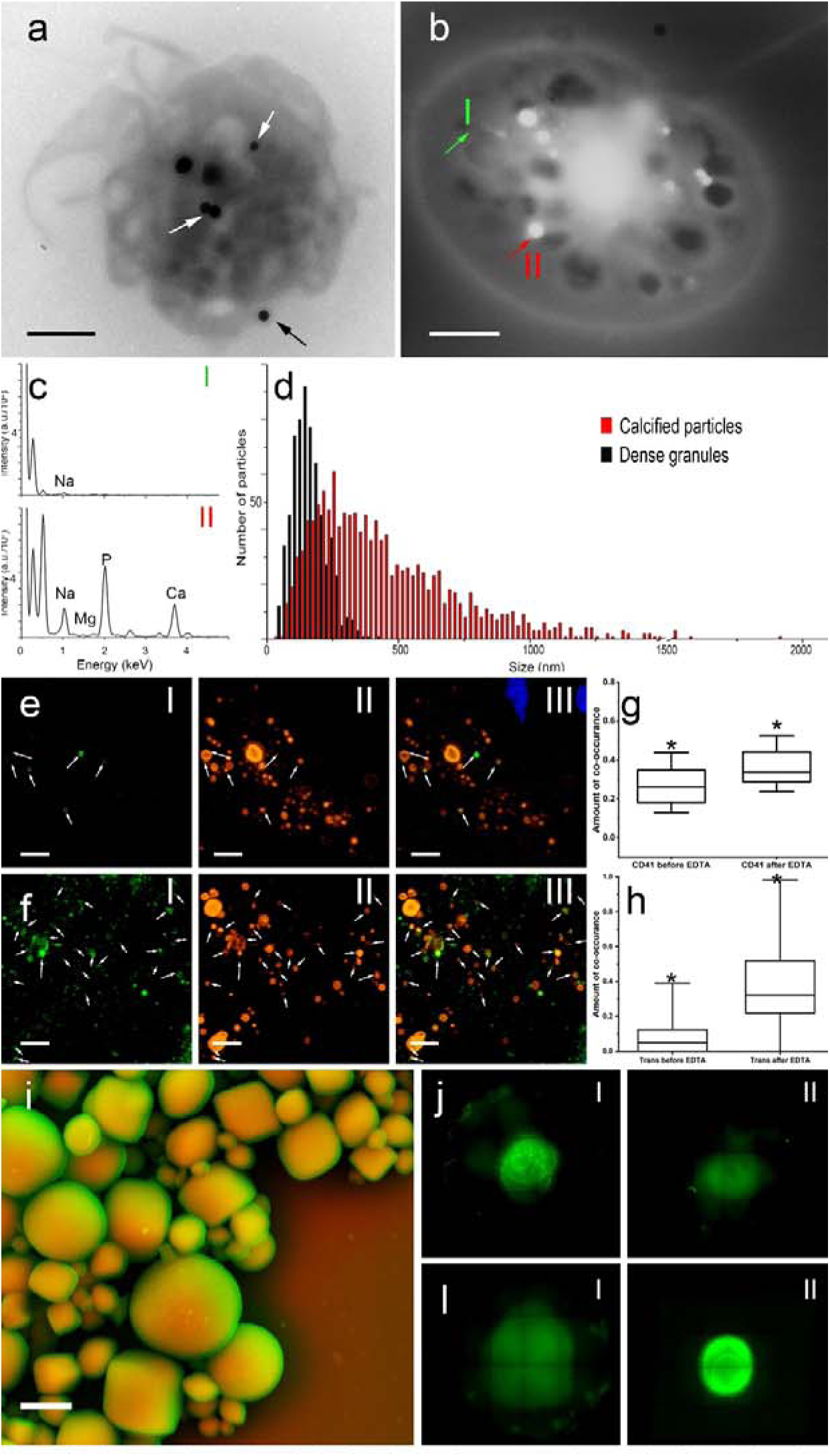
Electron microscopy analysis of a platelet, fluorescence micrographs of aortic tissue including: calcified particles, CD41, serotonin transporter, isolated calcified particles, and dot blot of proteins isolated from those particles. **a** TEM of a platelet, showing δ-granules highlighted by arrows. Scale bar = 1 µm. **b** Scanning-TEM of a platelet, where δ-granules are indicated by a green arrow. Scale bar = 1 µm. **c** EDS spectra of a platelet (**I**) and δ-granules (red arrow, **II**) of the platelet presented in **b**. **d** Size distribution of platelet δ-granules and calcified particles. **e** Fluorescence micrograph of histological slide of aorta showing CD41 staining (**I**), calcified particles stained by OsteoSense (**II**), and overlay micrographs, with nuclei in blue (**III**). Arrows indicate the regions presenting both CD41 and calcified particles. Scale bar = 10 µm. **f** Fluorescence micrograph of histological slide of aorta showing serotonin transporter staining (**I**), calcified particles stained by OsteoSense (**II**), and overlay micrographs, with nuclei in blue (**III**). Arrows indicate the regions presenting both the serotonin transporter and calcified particles. Scale bar = 10 µm. **g** Co-localization of CD41 with OsteoSense staining of aorta, with and without EDTA treatment. p = 0.04128. **h** Co-localization of serotonin transporter with OsteoSense staining of aorta, with and without EDTA treatment. p < 0.0001. **i** DDC-SEM of calcified particles isolated from aorta. Scale bar = 1 µm. **j** Dot blot from platelets stained with CD41 (**I**) and serotonin transporter (**II**). **l** Dot blot from proteins isolated from calcified particles stained with CD41 (**I**) and serotonin transporter (**II**).

To test this hypothesis, we firstly looked for platelet markers in the calcified particles, using immunofluorescence. Histological sections of aortas were stained for platelet markers CD41 (one of the few exclusive platelet markers^41^), serotonin transporter (which is highly expressed in platelets^42^ and δ-granules^43,44^), and calcium phosphate. The fluorescence micrographs confirm the presence of calcified particles, the platelet marker CD41 and serotonin transporter within the tissue (Fig. 2eI and 2fI). The micrographs also show (Fig. 2eI and 2fI) initially some degree of overlap between the platelet markers and the calcified particles (Fig. 2eIII and 2fIII), supporting previous reports in the literature where the same CD41 markers are observed in the calcification of human atherosclerotic plaques^45^.

Supposing that some of the structures presenting platelet markers are δ-granules trapped in the vasculature, and that some are calcified δ-granules, where the calcification would hinder access to the antigens, we attempted to slightly dissolve the calcification (mineral component), which might further expose any epitopes. To that end, an ethylenediaminetetraacetic acid (EDTA) solution was placed over new histological slides for a short period prior to the fluorescence staining. EDTA is a strong calcium chelating agent, and therefore its presence would slightly dissolve the calcium phosphate on the calcified particles, exposing epitopes of any platelet markers that might be present. New fluorescence micrographs were taken from the samples treated with EDTA. These presented a significant increase in the overlapping of platelet markers and calcification (Fig. 2g and h), suggesting again that platelets markers are indeed present in the calcified particles. To further confirm the presence of platelet markers in the calcified particles, particles isolated from aorta tissue (Fig. 2i) were subject to demineralization and protein extraction. The proteins extracted were evaluated by dot blot, and these were indeed positive for CD41 and serotonin transporter (Fig. 2l, 2m and Supplementary Figure S1).

Our next step was to test whether platelets can produce calcified particles *in vitro*. We developed an *in vitro* model to simulate the biological environment of the vasculature ECM where platelets and δ-granules would be trapped. The model consisted in having high density collagen gels (which have been commonly used as an *in vitro* simulator to vascular tissue^14^) with platelets incubated in serum (Fig. 3a). After 14 days, the gels were stained with DAPI for δ-granule visualization (the complex DAPI-polyphosphates formed within δ-granules can be excited at 480nm with an emission peak at 520 nm^46^) and OsteoSense for calcification visualization (Fig. 3b). It is clear from the images we obtained that calcified particles appeared in the gels presenting platelets (Fig 3bI). On the other hand, no calcified particle could be detected in gels incubated just with serum, even after 28 days of incubation (Fig 3bII). SEM micrographs and EDS analysis of the gels presenting platelets confirmed the presence of calcified particles (Fig. 3c) that presented the same morphology, size (around 200nm) and elemental composition as the particles found in the human aorta samples described earlier (Fig. 1c, 1d, 2d, and 2e) and reported in the literature^13^.

**Figure 3:**
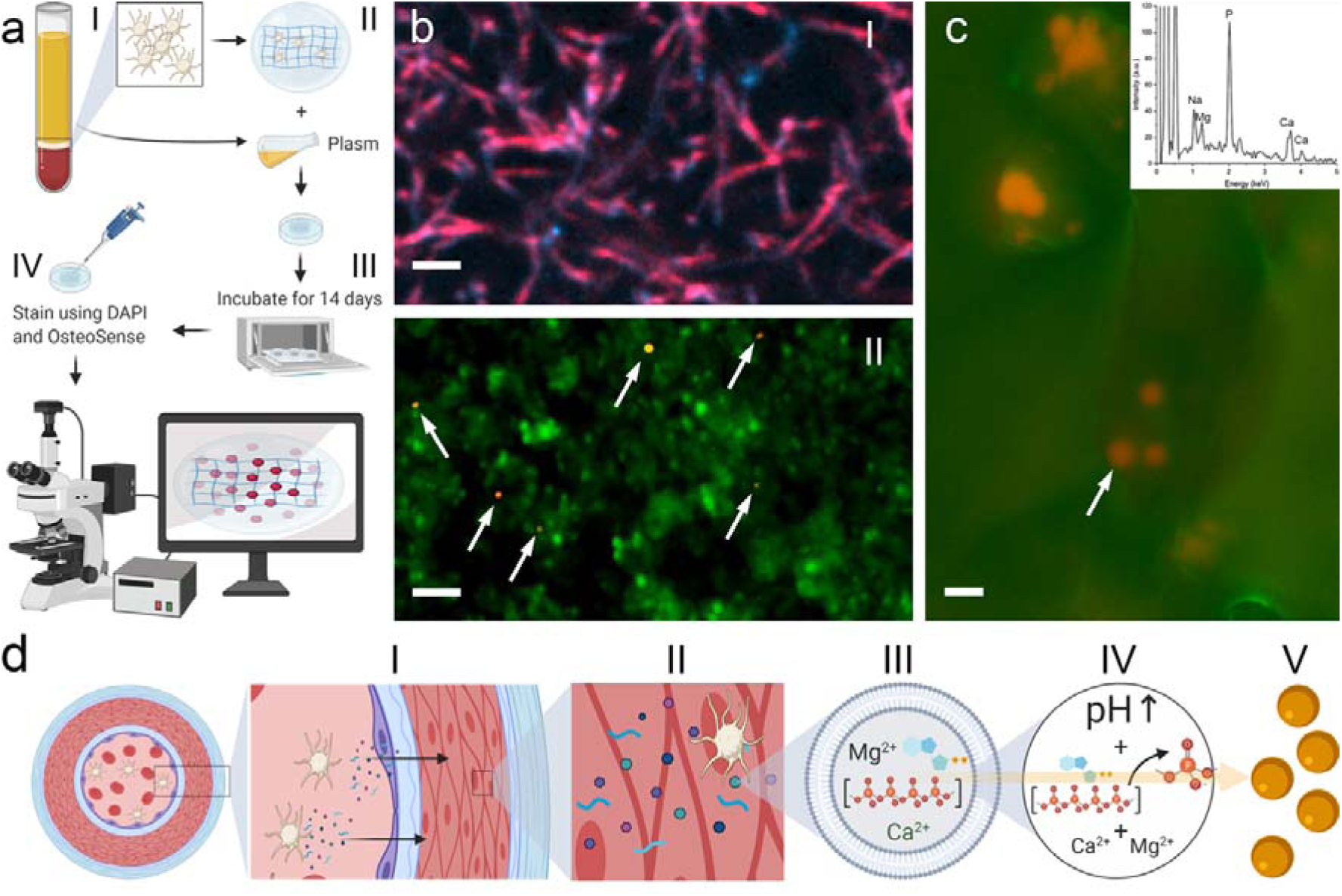
*In vitro* calcification model reproducing the calcified particles found *in vivo* and proposed mechanism of formation of calcified particles. **a** Diagram depicting the *in vitro* procedure designed to reproduce the calcified particles found *in vivo*. **I** Isolated platelets are mixed with collagen **II**. The solution is converted into a gel and then added to a serum free from platelets. **III** The gel is incubated for 14 days at 37 °C. **IV** After 14 days, the gel is stained with Osteosense and Dapi, then imaged by SEM and fluorescence microscopy. **b** Fluorescence micrograph of gel incubated for 14 days with serum free from platelets (**I**, collagen fibres as red and blue), presenting collagen fibres and gel containing platelets (**II**, platelets as green staining with δ-granules as brighter spots) and calcified particles (orange, arrows). Scale bar = 2 μm. **c** DDC-SEM micrograph and EDS of gel with platelets after 14 days in serum. Scale bar = 1 µm. **d** Schematic of proposed mechanism of formation of calcified particles found in vascular tissue. **I** Platelets release granules in the blood stream, and among these, δ-granules. **II** Platelet granules and platelets infiltrate artery walls and some of the granules, platelets and δ-granules are trapped in the tissue. **III** δ-granules contain within their interior calcium, magnesium, polyphosphate and adenosine di-phosphate (ADP)^37^, and when they are inside platelets have pH value = 5.4. **IV** When trapped in the tissue, their pH value increases to physiological pH (7.4), as their polyphosphate and ADP are degraded into ionic phosphate. **V** The increase in pH, combined with the formation of ionic phosphate, ionic calcium, and magnesium precipitates whitlockite inside the δ-granules, ultimately forming the calcified particles observed in the tissue.

Taking all of our results into account, we propose a mechanism of formation of calcified spherical particles (Fig. 3d). Initially, the natural activation of platelets would result in the release of δ-granules (Fig. 3dI). The δ-granules (or even some platelets), would then infiltrate the vascular wall (Fig. 3dI and II, as previously reported^26,47^) and get trapped in the ECM of the vasculature. Over time, the polyphosphate and Adenosine Diphosphate (ADP)^34,36^ within the interior of the δ-granules would dissociate^46^ (Fig. 3dIII and 3dIV), releasing ionic phosphate within the δ-granules. This ionic phosphate would then combine with magnesium and calcium (present in the higher concentration of 2.2M^35^) and following the resulting increase of pH to 7.4 to balance with the pH of ECM in the vasculature (initial value was 5.4, inside the δ-granules^48^), whitlockite would precipitate within the δ-granules (Fig. 3dV). The spherical shape of the δ-granules would be serving as a template/scaffold for the formation of the calcified particles, providing their size and spherical morphology.

Indeed, the formation of calcified particles would happen routinely on the tissue, even without any sign of cardiovascular disease or calcific lesions. This model would explain the presence of calcified particles on vascular tissues from humans at early ages and presenting no calcific disease, as has been reported in the literature^49^.

To confirm the hypothesis that the formation of the calcified particles is possibly routinely happening in vascular tissue, we searched for calcified particles on digested tissue from 48 healthy organ donors aged from 9 months to 78 years old (Supplementary info Table S1). Indeed, we could detect the presence of these particles in every single sample evaluated (Fig. 4a). Taking our hypothesis further, we would expect to find these particles in the vasculature of any animal presenting δ-granules in their thrombocytes. We looked for these particles on the vasculature of several mammals, birds and lizards (Supplementary info Table S2 for animal information) and we observed calcified particles in all of these classes (Fig. 4b, c, d, e and Supplementary Fig. S2). Indeed, the calcified particles observed in all these animals reproduce not only morphology and size, but also elemental composition, crystallinity and material phase (formed from whitlockite - Fig. 4f, g and h) of the particles reported in human vasculature.

**Figure 4:**
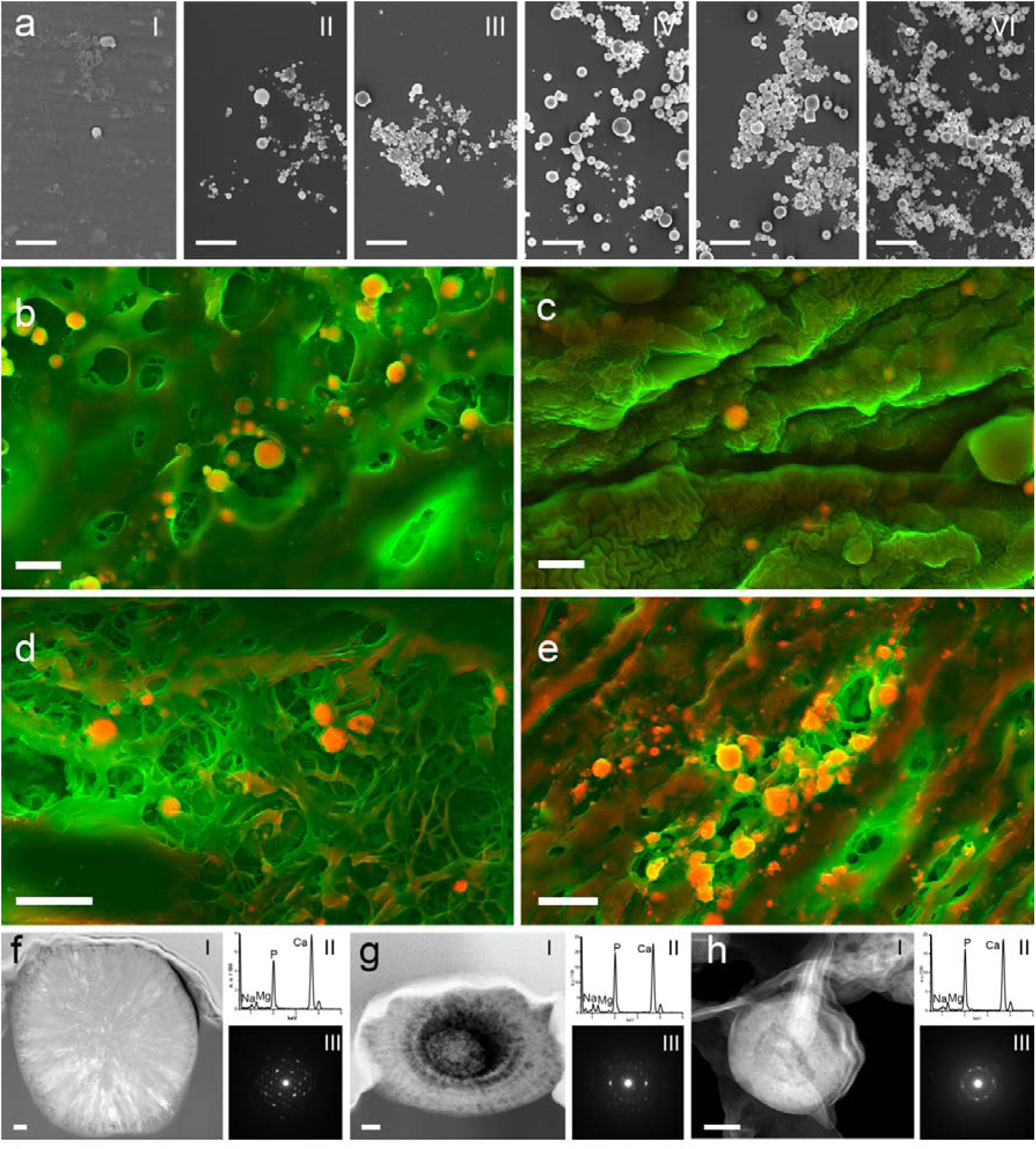
Occurrence, composition and crystallinity of calcified particles in vascular tissues from organ donors at different ages and from different classes of vertebrates (mammals, birds and lizards). **a** Representative scanning electron micrographs of calcified particles isolated from human aortic tissues obtained from individuals at (**I**) 9 months old, (**II**) 17 years old, (**III**) 31 years old, (**IV**) 38 years old, (**V**) 54 years old and (**VI**) 61 years old. Scale bars = 5 µm. DDC-SEM micrographs of vascular tissues from **b**, a rhesus macaque; **c**, a dog; **d**, a cassowary; **e**, an iguana. Scale bars = 2 µm. **f, g** and **h**, TEM (I), energy dispersive X-ray spectroscopy traces (**II**), and selected area electron diffraction patterns (**III**) of focused ion beam-sectioned calcified particles isolated from **c**, **d,** and **e** respectively. Scale bars = 100 nm.

Taking into account the ubiquity of these calcified particles, our model for the origin and formation of the calcified particles could also suggest the presence of the same or similar calcified particles in diseases where platelets or platelet particles (δ-granules) can be present.

Indeed, the literature reports the presence of similar particles (same morphology, size, elemental composition, crystallinity and material phase) as a consequence of different calcific diseases affecting tissues such as the eye^50^, cartilage^51^, and tumours^52^, with all the diseases associated to these tissues being strongly linked to platelets or platelet particles^41,53–55^. In summary, the results presented here may not only have implications for the understanding of cardiovascular calcific diseases, but could also have implications for other diseases where calcification is present.

On the other hand, our newly proposed formation mechanism and the origins of the spherical calcified particles described herein suggest that calcification (in particular the vascular calcification) may involve multiple aetiologies. It is essential to emphasize that our model does not invalidate existing mechanisms proposed in the literature. Rather, our findings complement models describing subsequent calcification processes (occurring both after and parallel to the formation of δ-granule-derived calcified particles), wherein the propagation of calcified tissue, mediated by cardiac or other cell types, leads to the development of calcified fibers and compact calcification^13^.

The mechanisms described in the existing literature would be strongly related to cardiovascular diseases and their biological and cellular mechanisms^14,56^. This existing literature shows that this secondary calcification is strongly associated with osteogenic processes and inflammation marked by the presence of neutrophils^57^. To explore this possible correlation to osteogenicity and inflammation, we stained the healthy aortic tissue with osteopontin, Runx2 (osteogenic markers), and a neutrophil marker (CD11).

The micrographs confirm the presence of specific osteogenic markers (Fig. 5a) and neutrophils (Fig. 5b) in the vascular tissue from a healthy organ donor; however, these are not present in high densities. Furthermore, only a limited number of these markers are associated with calcified deposits. This observation aligns with the previous findings indicating that osteogenic markers are not associated with the calcified particles themselves^13^, further supporting the hypothesis that this early-stage calcification (spherical particles) originates from platelets rather than through osteogenic or inflammatory mechanisms.

**Figure 5:**
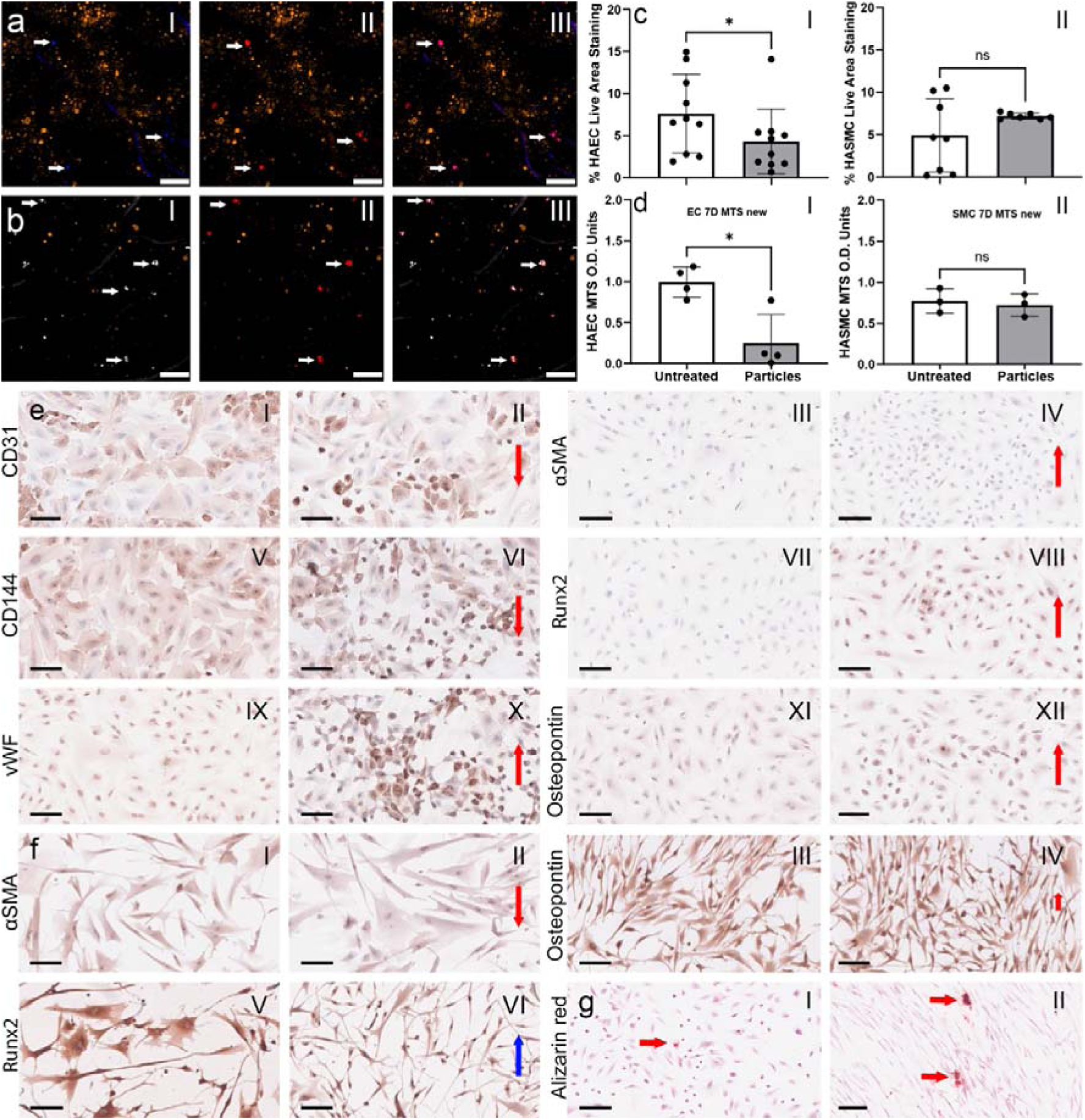
Fluorescence micrographs of aortic tissue analysing osteogenic and neutrophil signals, and live/dead and proliferation assay and protein expression of vascular cell culture with calcified particles isolated from human aorta. **a** Fluorescence micrograph of histological slide of aorta showing weak Runx2 staining (**I** - blue), weak osteopontin staining(**II** - red), and overlay micrographs (**III**). Calcified particles are stained with Osteosense (orange). Arrows indicate the regions presenting both Runx2 and osteopontin. Scale bar = 10 µm. **b** Fluorescence micrograph of histological slide of aorta showing CD11b (**I** - white), osteopontin (**II** - red), and overlay micrographs (**III**). Calcified particles are stained with Osteosense (orange). Arrows indicate the regions presenting both CD11b and osteopontin. Scale bar = 10 µm. **c** Live staining quantification for HAECs and HASMCs culture with calcified particles for 14 days (dead stain presented in Supplementary Fig. S3). **d** Proliferation assay (MTS) with HAECs and HASMCs for 7 days. **e** Protein expression for HAECs cultured without (**I**, **III**, **V, VI and IX**) and treated with calcified particles (**II**, **IV**, **VI, VIII and X**). Arrow up represents the upregulation of proteins, and arrow down represents the downregulation of proteins. **f** Protein expression for HASMCs cultured without (**I**, **III**, **V**) and treated with calcified particles (**II**, **IV**, **VI**). Arrow up represents the upregulation of proteins, and arrow down represents the downregulation of proteins, with the blue arrow representing upregulation in the nucleus. **g** Calcification assay (staining with alizarin red) with HAECs (**I**) and HASMCs (**II**) (controls presented at Supplementary Fig. S4 with also Von Kossa staining showed no positivity for alizarin red).

Finally, a critical question remains regarding the functional role of these platelet-derived calcified particles in cardiovascular pathology. Specifically, whether they interact with and influence vascular cells to promote calcification, and whether they could play a role in the early stages of cardiovascular calcific disease. These particles may interact with vascular cells (consistent with previous observations involving synthetic calcium phosphate^58^), thereby promoting cell differentiation and mineral deposition^59,60^.

To evaluate the possible impact of these particles on vascular biology, the two main types of vascular cells (human aortic smooth muscle cells (HASMCs) and human aortic endothelial cells (HAECs)) were cultured in the absence, and presence of calcified particles isolated from human aortas. On live/dead assay and proliferation assessment (Fig. 5c and Supplementary Fig. S3), it is clear that the particles have a significant deleterious effect on the HAECs (Fig. 5cI and 5dI) while presenting no effect on the HASMCs (Fig. 5cII and 5dII). These results are not surprising, considering that the particles are present and likely in contact with HASMCs throughout the entire life of the animal (as suggested in Figs. 1 and 4). Regarding the possible differentiation and osteogenic potential of the calcified particles, it is clear that they can induce vascular cells to an osteogenic phenotype. HAECs downregulate endothelial markers such as CD31 (Fig. 5eI and 5eII) and CD144 (Fig. 5eVII and 5eVIII) and upregulate αSMA(Fig. 5eIII and 5eIV), Runx2 (Fig. 5eV and 5eVI), vWF (Fig. 5eIX and 5eX), and osteopontin (Fig. 5eXI and 5eXII). vWF is upregulated in response to injury, stress and pro-thrombotic signals and mediates platelet adhesion. Upregulation of αSMA indicates the onset of endothelial to mesenchymal transition (EMT). Finally, the upregulation of Runx2 and osteopontin suggests a transition to osteogenic phenotype. A similar pattern suggesting a transition to osteogenic phenotype is observed for the HASMCs (Fig. 5f). Here αSMA is downregulated (Fig. 5fI and 5fII), a small upregulation for osteopontin (Fig. 5fIII and 5fIV), and an upregulation in the nuclei (suggesting the activation of the protein) for Runx2 (Fig. 5fV and 5fVI).

The osteogenic potential is further supported by the observation that cells begin to produce calcified nodules, with HASMCs (Fig. 5gII) showing a slightly stronger effect than HAECs (Fig.5gI, (controls presented at Supplementary Fig. S4 with also Von Kossa staining). Taken together, these results suggest that the calcified particles are not merely a passive component of the tissue, but rather one can actively influence vascular cells, inducing them to differentiate and upregulate osteogenic-related proteins and even promoting mineral deposition (Fig. 5g). It is important to highlight that the cell cultures were maintained for 3 weeks with a limited number of calcified particles, and even within this short timeframe and with a limited particle load, vascular cells already exhibited an osteogenic response. One could speculate that the progressive accumulation of calcified particles within the tissue (which naturally occurs with aging) and prolonged exposure to these particles may, at some point during the animal’s life, constitute a trigger for the promotion of pathological calcification through their effects on vascular cells.

If confirmed, the proposed mechanism of formation and origins of calcified spherical particles from δ-granules and their influence on cardiac cells offers a glimpse of a potential pharmacological intervention in the development and treatment of cardiovascular calcific diseases. Until now, every pharmacological intervention designed either for the prevention or treatment of cardiovascular calcific diseases such as atherosclerosis, aortic valve stenosis, and rheumatic fever, has been unsuccessful. Pharmacological interventions that could interfere with the number of δ-granules being trapped in the vasculature ECM or interfere on the influence of the calcified particles on vascular cells could ultimately prevent the accumulation of calcified particles in the tissue and the triggering of other mineralization mechanisms. As a consequence, such interventions could aid in mitigating the damaging effects that these particles would have in the tissue, and either slow down or stop the progression of cardiovascular calcific disease.

## Acknowledgements

The authors would like to acknowledge and to thank Magdalena Unterberger, Saumya Singh, and Joseph A. M. Steele for their help with the development and initial tests for fluorescence and embedding and sectioning of tissues. We gratefully acknowledge the role of the Oxford Cell and Tissue Biobank in collecting and making samples and/or data available. We are also grateful to the patients who generously donated their tissues and shared their data to be used in the generation of this publication. We would like to thank Prof. Susan Evans for providing cardiac tissue from reptiles, and Dr Charlotte Brassey for providing cassowary cardiac tissue. Schematics in Figure 1 and 3 were created using Biorender.com.

## Author Contributions

S.B. and E.T. performed sample preparation, conducted electron and fluorescence microscopy work and performed gel experiments. S.B. and E.T. did blotting experiments. S.A. did FIB sectioning and electron diffraction and aided with data interpretation. K.S., J.T., M.H.Y and A.H.C. helped with human tissue sample procurement and I.K.H. aided in data interpretation. N.L., A.M., P.S., did the cell culture and helped with manuscript writing and data interpretation. S.B. conceived, designed and coordinated the study, interpreted the data and wrote the manuscript.

## References

1 Mc Namara, K., Alzubaidi, H. & Jackson, J. K. Cardiovascular disease as a leading cause of death: how are pharmacists getting involved? Integrated pharmacy research & practice 8, 1–11 (2019).

2 Tölle, M., Reshetnik, A., Schuchardt, M., Höhne, M. & van der Giet, M. Arteriosclerosis and vascular calcification: causes, clinical assessment and therapy. European journal of clinical investigation 45, 976–985 (2015).

3 Peeters, F. et al. Calcific aortic valve stenosis: hard disease in the heart: A biomolecular approach towards diagnosis and treatment. Eur Heart J 39, 2618–2624 (2018).

4 Leal, M. et al. Rheumatic heart disease in the modern era: recent developments and current challenges. Revista da Sociedade Brasileira de Medicina Tropical 52, e20180041 (2019).

5 Demer, L. L. & Tintut, Y. Vascular Calcification: Pathobiology of a Multifaceted Disease. Circulation 117, 2938–2948 (2008).

6 Zazzeroni, L., Faggioli, G. & Pasquinelli, G. Mechanisms of Arterial Calcification: The Role of Matrix Vesicles. European Journal of Vascular and Endovascular Surgery 55, 425–432 (2018).

7 Durham, A. L., Speer, M. Y., Scatena, M., Giachelli, C. M. & Shanahan, C. M. Role of smooth muscle cells in vascular calcification: implications in atherosclerosis and arterial stiffness. Cardiovasc Res 114, 590–600 (2018).

8 Tintut, Y., Hsu, J. J. & Demer, L. L. Lipoproteins in Cardiovascular Calcification: Potential Targets and Challenges. Frontiers in cardiovascular medicine 5, 172–172 (2018).

9 Kapustin, A. N. et al. Vascular Smooth Muscle Cell Calcification Is Mediated by Regulated Exosome Secretion. 116, 1312–1323 (2015).

10 Xie, C. et al. The Emerging Role of Mesenchymal Stem Cells in Vascular Calcification. Stem Cells International 2019, 2875189 (2019).

11 Ho, C. Y. & Shanahan, C. M. Medial Arterial Calcification: An Overlooked Player in Peripheral Arterial Disease. Arterioscler Thromb Vasc Biol 36, 1475–1482 (2016).

12 Hsu, J. J., Lim, J., Tintut, Y. & Demer, L. L. Cell-matrix mechanics and pattern formation in inflammatory cardiovascular calcification. Heart 102, 1710–1715 (2016).

13 Bertazzo, S. et al. Nano-analytical electron microscopy reveals fundamental insights into human cardiovascular tissue calcification. Nat Mater 12, 576–583 (2013).

14 Hutcheson, J. D. et al. Genesis and growth of extracellular-vesicle-derived microcalcification in atherosclerotic plaques. Nat. Mater. 15, 335 (2016).

15 New, S. E. et al. Macrophage-derived matrix vesicles: an alternative novel mechanism for microcalcification in atherosclerotic plaques. Circ Res 113, 72–77 (2013).

16 Reynolds, J. L. et al. Human vascular smooth muscle cells undergo vesicle-mediated calcification in response to changes in extracellular calcium and phosphate concentrations: a potential mechanism for accelerated vascular calcification in ESRD. J Am Soc Nephrol 15, 2857–2867 (2004).

17 Mohr, W. & Gorz, E. Does calcification of arteriosclerotic vessels imitate osteogenesis? Pathomorphological study of arteriosclerotic plaques. Zeitschrift für Kardiologie 91, 212-+ (2002).

18 Demer, L. L. & Abedin, M. Skeleton key to vascular disease. J Am Coll Cardiol 44, 1977–1979 (2004).

19 Burgstahler, C. et al. Elevated coronary calcium scores are associated with higher residual platelet aggregation after clopidogrel treatment in patients with stable angina pectoris. International Journal of Cardiology 135, 132–135 (2009).

20 Jayachandran, M. et al. Characterization of blood borne microparticles as markers of premature coronary calcification in newly menopausal women. Am. J. Physiol.-Heart Circul. Physiol. 295, H931–H938 (2008).

21 Lv, H. et al. Doxorubicin contributes to thrombus formation and vascular injury by interfering with platelet function. Am J Physiol Heart Circ Physiol 319, H133–h143 (2020).

22 Liu, O. et al. Clopidogrel, a Platelet P2Y12 Receptor Inhibitor, Reduces Vascular Inflammation and Angiotensin II Induced-Abdominal Aortic Aneurysm Progression. PLoS One 7, e51707 (2012).

23 Diehl, P. et al. Increased levels of circulating microparticles in patients with severe aortic valve stenosis. Thromb Haemost 99, 711–719 (2008).

24 Puhm, F., Boilard, E. & Machlus, K. R. Platelet Extracellular Vesicles: Beyond the Blood. Arterioscler Thromb Vasc Biol 41, 87–96 (2021).

25 Farr, M., Wainwright, A., Salmon, M., Hollywell, C. A. & Bacon, P. A. Platelets in the synovial fluid of patients with rheumatoid arthritis. Rheumatol. Int. 4, 13–17 (1984).

26 Ho-Tin-Noé, B., Boulaftali, Y. & Camerer, E. Platelets and vascular integrity: how platelets prevent bleeding in inflammation. Blood 131, 277–288 (2018).

27 Boilard, E. et al. Platelets amplify inflammation in arthritis via collagen-dependent microparticle production. Science 327, 580–583 (2010).

28 French, S. L. et al. Platelet-derived extracellular vesicles infiltrate and modify the bone marrow during inflammation. Blood Advances 4, 3011–3023 (2020).

29 Boilard, E., Duchez, A. C. & Brisson, A. The diversity of platelet microparticles. Curr Opin Hematol 22, 437–444 (2015).

30 van der Meijden, P. E. J. & Heemskerk, J. W. M. Platelet biology and functions: new concepts and clinical perspectives. Nature Reviews Cardiology 16, 166–179 (2019).

31 Burnouf, T. et al. An overview of the role of microparticles/microvesicles in blood components: Are they clinically beneficial or harmful? Transfus Apher Sci 53, 137–145 (2015).

32 Manne, B. K., Xiang, S. C. & Rondina, M. T. Platelet secretion in inflammatory and infectious diseases. Platelets 28, 155–164 (2017).

33 Meyers, K. M., Holmsen, H. & Seachord, C. L. Comparative study of platelet dense granule constituents. The American journal of physiology 243, R454–461 (1982).

34 Chen, Y., Yuan, Y. & Li, W. Sorting machineries: how platelet-dense granules differ from α-granules. Biosci Rep 38, BSR20180458 (2018).

35 Holmsen, H. & Weiss, H. J. Secretable storage pools in platelets. Annual review of medicine 30, 119–134 (1979).

36 Ruiz, F. A., Lea, C. R., Oldfield, E. & Docampo, R. Human platelet dense granules contain polyphosphate and are similar to acidocalcisomes of bacteria and unicellular eukaryotes. The Journal of biological chemistry 279, 44250–44257 (2004).

37 Ruiz, F. A., Lea, C. R., Oldfield, E. & Docampo, R. Human Platelet Dense Granules Contain Polyphosphate and Are Similar to Acidocalcisomes of Bacteria and Unicellular Eukaryotes. J. Biol. Chem. 279, 44250–44257 (2004).

38 Meyers, K. M., Holmsen, H. & Seachord, C. L. Comparative-study of platelet dense granule constituents. American Journal of Physiology 243, R454–R461 (1982).

39 Massy, Z. A. & Drüeke, T. B. Magnesium and outcomes in patients with chronic kidney disease: focus on vascular calcification, atherosclerosis and survival. Clinical Kidney Journal 5, i52–i61 (2012).

40 Rendu, F. & Brohard-Bohn, B. The platelet release reaction: granules’ constituents, secretion and functions. Platelets 12, 261–273 (2001).

41 Boilard, E. et al. Platelets Amplify Inflammation in Arthritis via Collagen-Dependent Microparticle Production. Science 327, 580–583 (2010).

42 Dmitriev, A. D. et al. Western blot analysis of human and rat serotonin transporter in platelets and brain using site-specific antibodies: evidence that transporter undergoes endoproteolytic cleavage. Clinica chimica acta; international journal of clinical chemistry 356, 76–94 (2005).

43 Ambrosio, A. L. & Di Pietro, S. M. Storage pool diseases illuminate platelet dense granule biogenesis. Platelets 28, 138–146 (2017).

44 Chatterjee, D. & Anderson, G. M. The human platelet dense granule: serotonin uptake, tetrabenazine binding, phospholipid and ganglioside profiles. Archives of biochemistry and biophysics 302, 439–446 (1993).

45 Foresta, C. et al. Platelets express and release osteocalcin and co-localize in human calcified atherosclerotic plaques. J. Thromb. Haemost. 11, 357–365 (2013).

46 Omelon, S. J. & Grynpas, M. D. Relationships between Polyphosphate Chemistry, Biochemistry and Apatite Biomineralization. Chem. Rev. 108, 4694–4715 (2008).

47 Boilard, E., Blanco, P. & Nigrovic, P. A. Platelets: active players in the pathogenesis of arthritis and SLE. Nat. Rev. Rheumatol. 8, 534–542 (2012).

48 Carty, S. E., Johnson, R. G. & Scarpa, A. Serotonin transport in isolated platelet granules. Coupling to the electrochemical proton gradient. J. Biol. Chem. 256, 11244–11250 (1981).

49 Eggen, D. A., Strong, J. P. & McGill, H. C., Jr. Coronary calcification. Relationship to clinically significant coronary lesions and race, sex, and topographic distribution. Circulation 32, 948–955 (1965).

50 Tan, A. C. et al. Calcified nodules in retinal drusen are associated with disease progression in age-related macular degeneration. Science translational medicine 10, eaat4544 (2018).

51 Scotchford, C. A., Vickers, M. & Ali, S. Y. THE ISOLATION AND CHARACTERIZATION OF MAGNESIUM WHITLOCKITE CRYSTALS FROM HUMAN ARTICULAR-CARTILAGE. Osteoarthritis and Cartilage 3, 79–94 (1995).

52. Tsolaki, E. et al. Invasive breast tumors are characterized by the presence of crystalline nanoparticles. 2020.2004.2029.067660 (2020).

53 Labelle, M., Begum, S. & Hynes, Richard O. Direct Signaling between Platelets and Cancer Cells Induces an Epithelial-Mesenchymal-Like Transition and Promotes Metastasis. Cancer cell 20, 576–590 (2011).

54 Gay, L. J. & Felding-Habermann, B. Contribution of platelets to tumour metastasis. Nature Reviews Cancer 11, 123–134 (2011).

55 Sengul, E. A. et al. Correlation of neutrophil/lymphocyte and platelet/lymphocyte ratio with visual acuity and macular thickness in age-related macular degeneration. Int J Ophthalmol 10, 754–759 (2017).

56 Kapustin, A. N. et al. Vascular Smooth Muscle Cell Calcification Is Mediated by Regulated Exosome Secretion. Circ.Res. 116, 1312–1323 (2015).

57 Lindman, B. et al. Calcific aortic stenosis. Nature Reviews Disease Primers 2 (2016).

58 Liu, Q. et al. Nano-hydroxyapatite accelerates vascular calcification via lysosome impairment and autophagy dysfunction in smooth muscle cells. Bioactive Materials 8, 478–493 (2022).

59 Liu, H., Huang, L.-H., Sun, X.-Y. & Ouyang, J.-M. High-phosphorus environment promotes calcification of A7R5 cells induced by hydroxyapatite nanoparticles. Materials Science and Engineering: C 107, 110228 (2020).

60 Yuan, H. et al. Osteoinductive ceramics as a synthetic alternative to autologous bone grafting. Proceedings of the National Academy of Sciences 107, 13614–13619 (2010).

