## Supplementary information for "Biomineralization from platelet δ-granules as the origin of cardiovascular calcification in humans and other animals"

**Table S1:** Patient demographics for human tissue samples. Aorta (n=48).

| **Age (yr)** | 4 days to 78 years |
| --- | --- |
| **Sex** | 44% male, 56% female |
| **Cause of death** | Subarachnoid haemorrhage 21%, Intracranial haemorrhage 13%, Hypoxic brain injury 13%, Ischemic heart disease 8%, Cardiac arrest 2%, Myocardial infarction 6%, Other 33%, unknown 4% |
| **Cause of rejection for valve transplant** | Micro-infected 52%, Failed/Missing virology test 10%, Calcified 21%, Atheroma 6%, Unknown10% |

**Table S2:** Animal vascular tissues.

| **Animal groups** | **Type** | **Kind / age** |
| --- | --- | --- |
| **Mammals** |  |  |
|  | **Wild-type** | 2 x piglet (8 weeks old and 4 days old)  1 x goat (4 years old)  1 x calf (5 months old)  1 x reindeer (3 years old and 3 months old)  3 x cats (5, 12 and 12.5 years old)  3 x dogs (Adult, 22 and 9 years old)  1 x sheep (2 years old)  1 x cria (2 days old)  1 x rabbit (5 years old)  1 x horse (6 years old)  1 x rat (0.5 year old)  1 x rhesus macaque (18 years old) |
| **Avian reptiles (birds)** |  | 1 x cassowary (-) |
| **Non-avian reptiles**  **(lizards, snakes, etc.)** |  | 1 x iguana (-) |

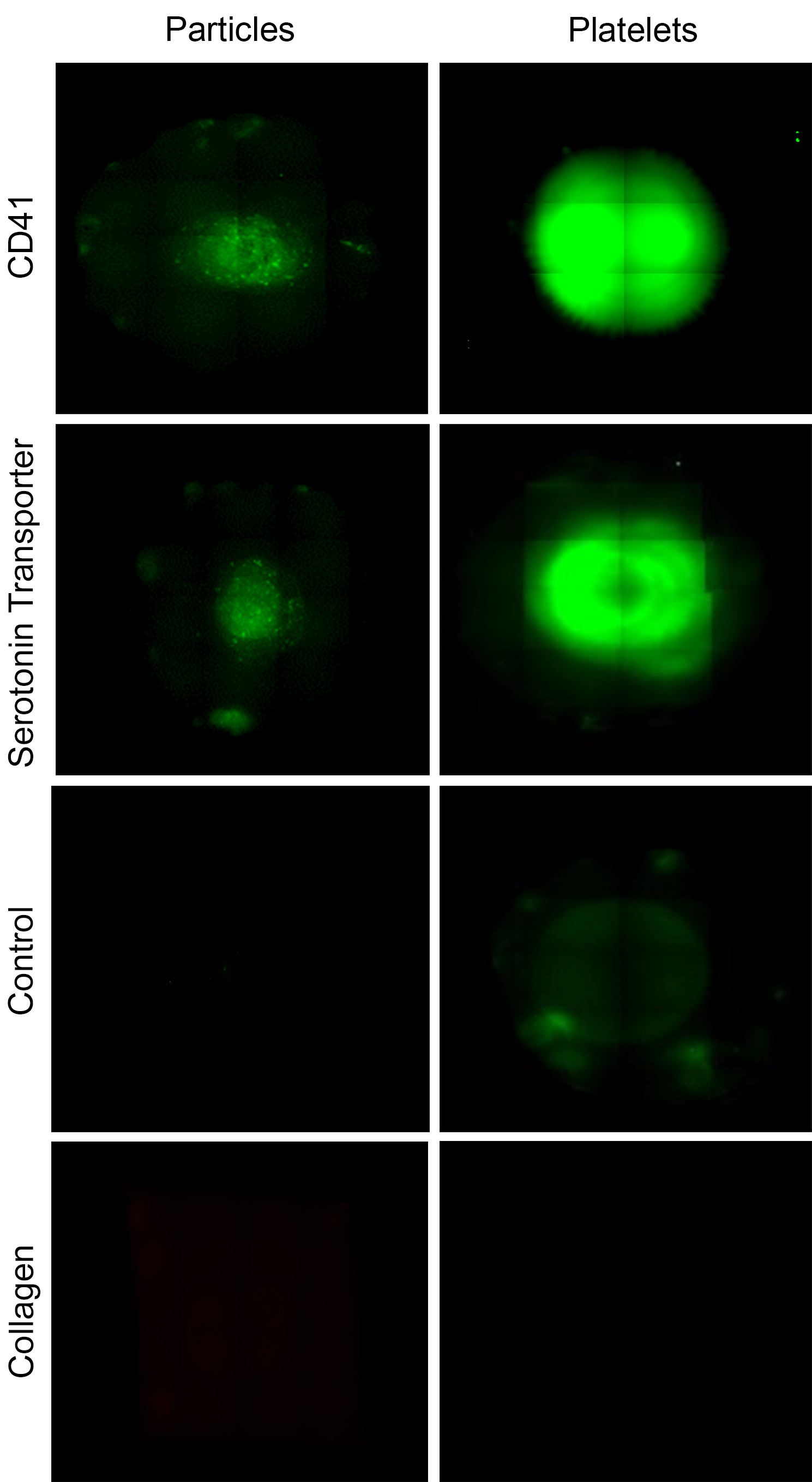

**Supplementary Figure 1: Dot blot of whitlockite nanoparticles and platelet lysates**. Positive staining was observed for CD41 and serotonin transporter antibodies. Controls omitting the primary antibodies indicated no staining. Both lysates were also stained for collagen where no staining was observed.

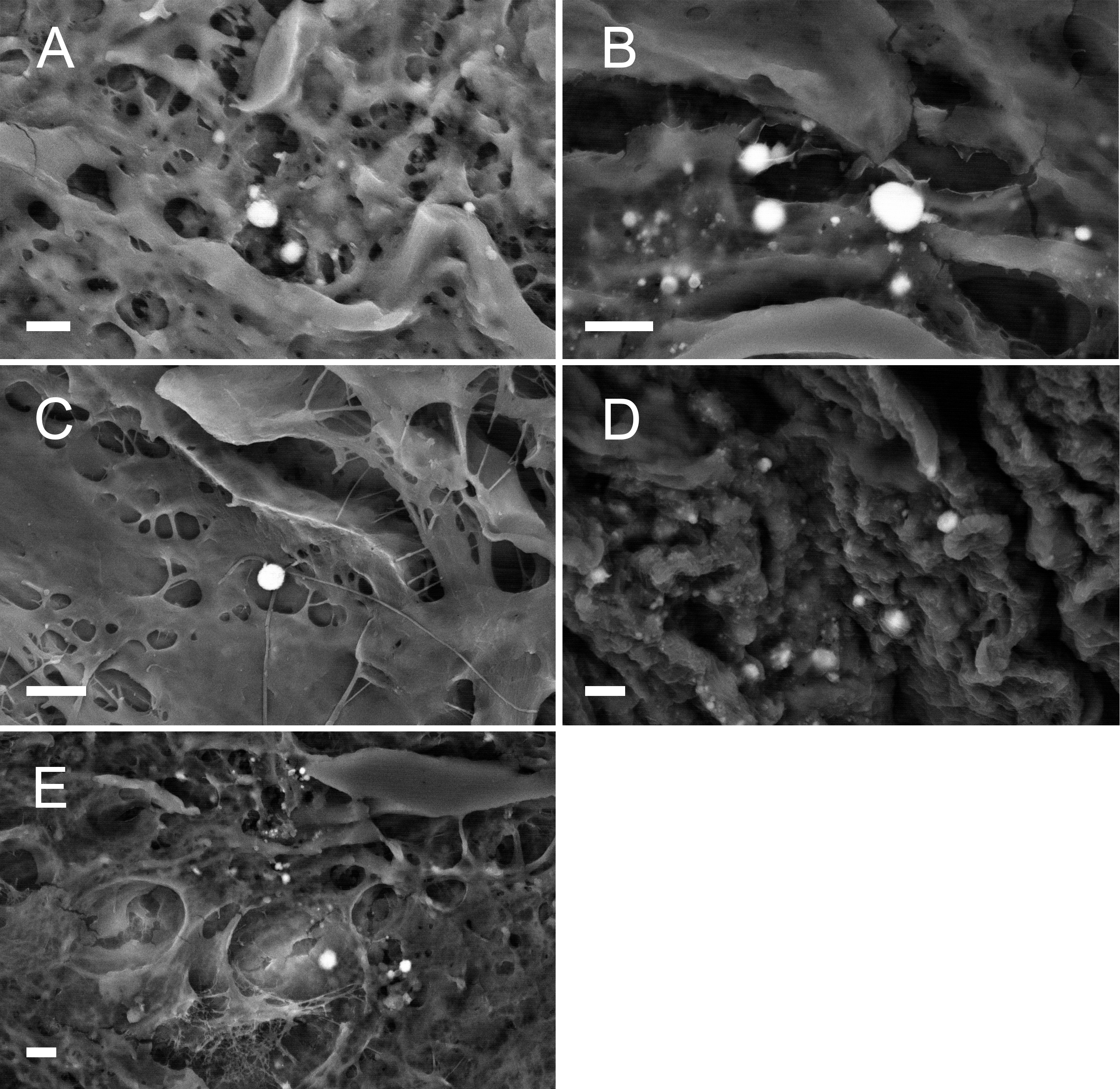

**Supplementary Figure 2:** SEM micrographs using backscattering detector of calcified particles in vascular tissue from (**A** and **B**) cats, (**C**) a goat, (**D**) a rabbit and (**E**) a horse. Scale bar = 2 µm.

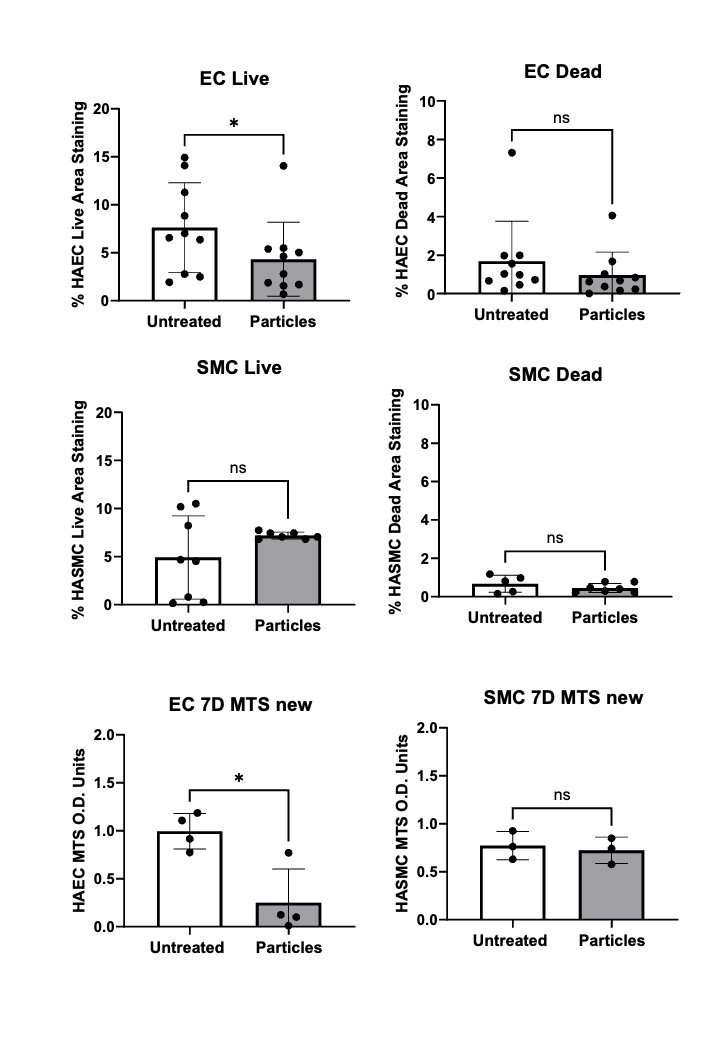

**Supplementary Figure 3: Live/dead and proliferation assay and protein expression of vascular cell culture with calcified particles isolated from human aorta.** Live staining quantification for HAECs and HASMCs culture with calcified particles for 14 days. Proliferation assay (MTS) with HAECs and HASMCs for 7 days.

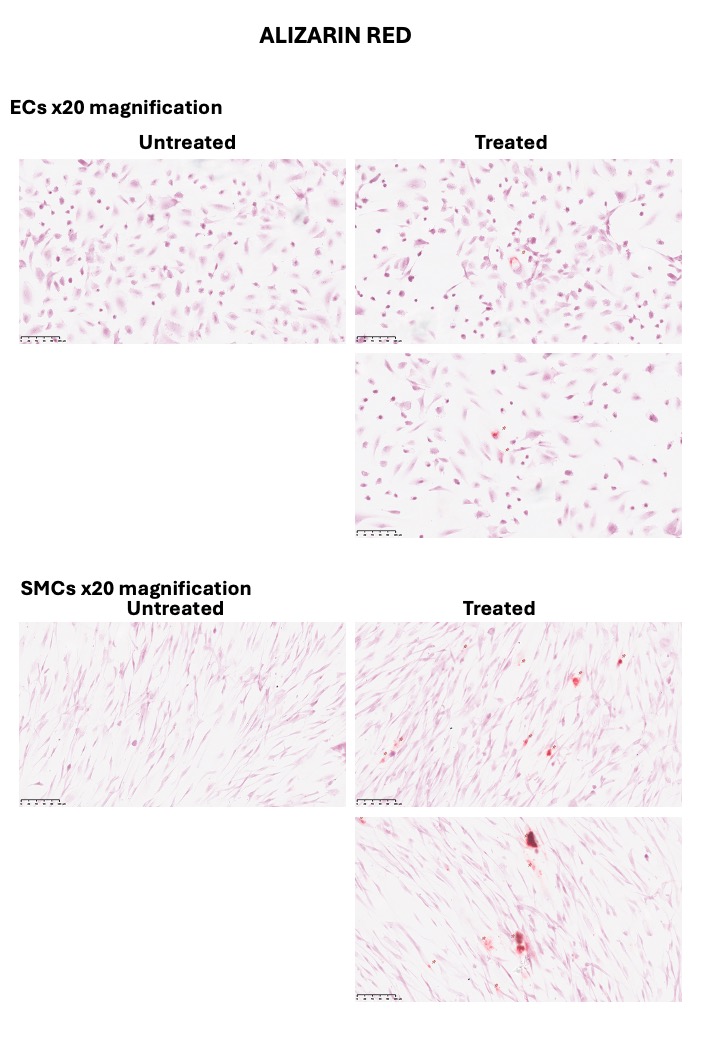

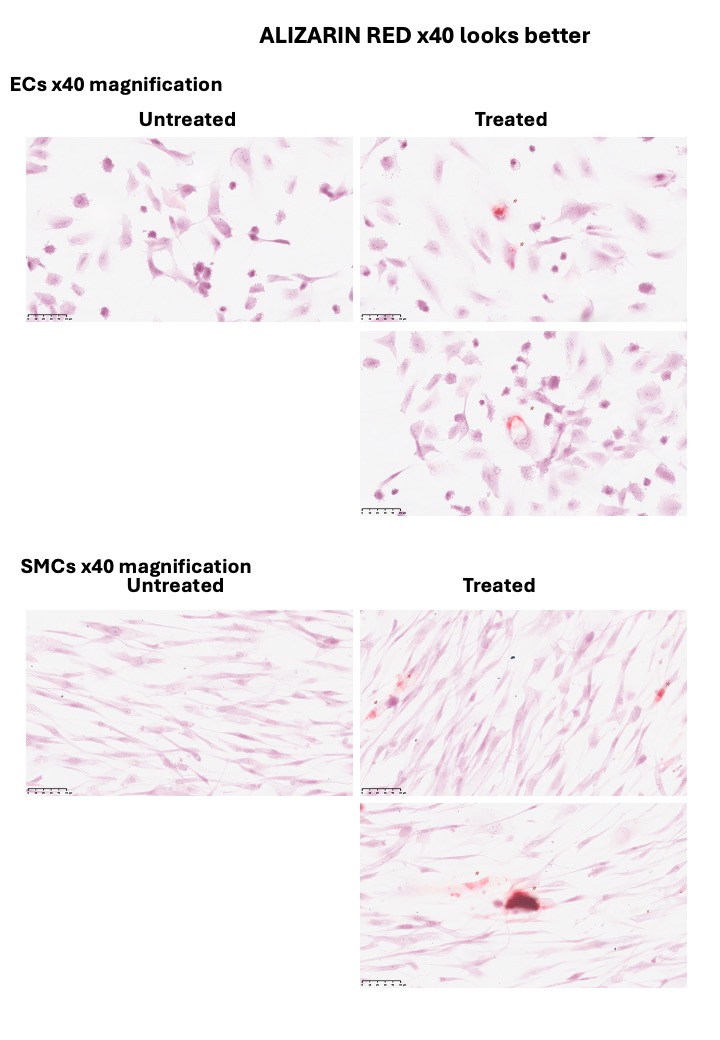

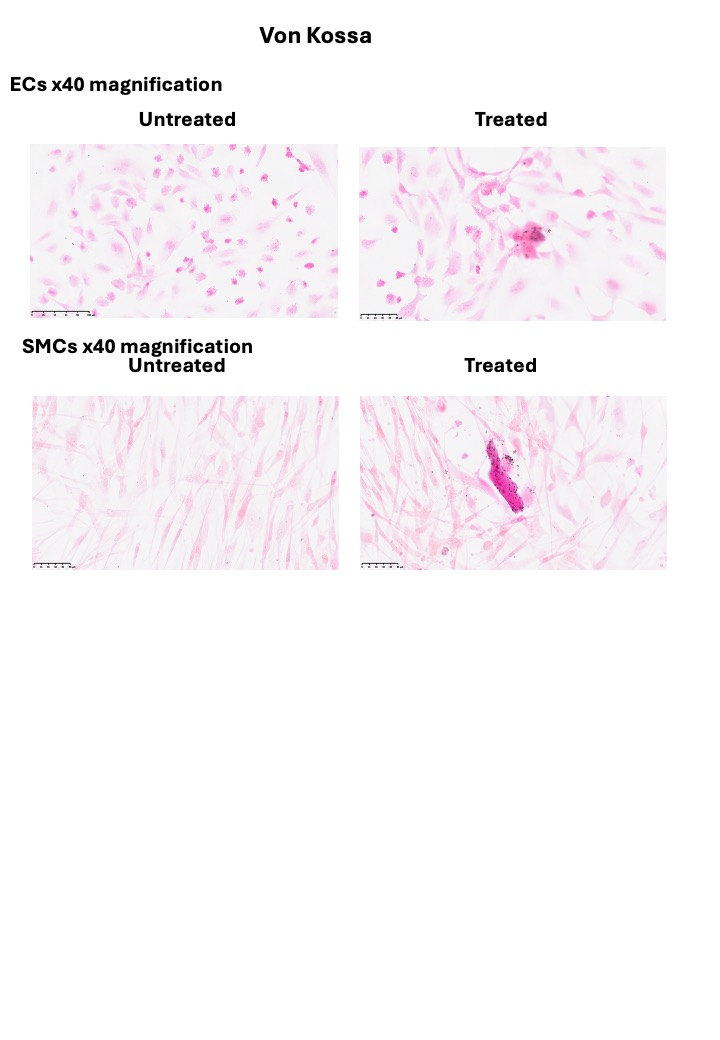

**Supplementary Figure 4:** Calcification assay staining with alizarin red or Von Kossa with HAECs and HASMCs treated with calcified particles.

**Methods**

***Human vascular tissues:*** Human tissues were obtained from the Oxford Heart Valve Bank (John Radcliffe Hospital, Oxford, UK) after they had been rejected for transplant. All samples were collected under approved ethical guidelines with informed consent that allowed the anonymous analysis of the tissues.

***Other vertebrate vascular tissues*:** Vascular tissue from vertebrates were provided by the Royal Veterinary College – London (Hertfordshire, UK), University College London (London, UK), National Museum of Scotland (Edinburgh, UK), Medical Research Council Centre for Macaques, Porton Down, Salisbury (Salisbury, UK). The samples were obtained from post-mortem autopsy of animals that had died of natural causes (see Table S2). The tissues were immediately fixed with 4% (w/v) formaldehyde (Sigma, BioReagent, ≥36.0%) solution in phosphate buffered saline (PBS; Sigma) at room temperature for at least 1 day.

***Platelet-rich plasma:*** Platelet-rich plasma was obtained in collaboration with Professor Janice Tsui from NHS blood and transplant biobank after a project ethical approval was obtained by the Proportionate Review Sub-Committee of the North East - Tyne & Wear South Research Ethics Committee (18/NE/-354).

***Histology and immunohistochemistry*:** Human aorta samples were embedded in paraffin wax, sectioned at 4 µm, and mounted on histological slides. Sections were dewaxed by being immersed twice in xylene and hydrated through an increased series of ethanol concentrations. Slides were stained with Harris haematoxylin and 1% (w/v) eosin Y (Sigma Aldrich, UK).

***Tissue digestion:*** The human aorta samples were incubated overnight at 37 ^o^C in 0.4% collagenase from Clostridium histolyticum, Type I (Sigma) solution in PBS. The tissues were then homogenized using a tapered tissue grinder. The tissue homogenate solutions were incubated one more time overnight at 37 ^o^C. Particles where then isolated using a sucrose gradient of saturated, 60% and 50% and centrifuged at 22000g. The particles were collected and washed with PBS.

***Florescence staining protocol:*** For this work, all samples to be immunofluorescence stained were fixed as quickly as possible following surgical removal using 10% formalin for at least 24 hours or were immediately processed to be cut into histological slides. Samples were then dehydrated and embedded in paraffin wax before being cut into 4 – *5* μm thick slices and placed on histological glass slides. Following that, samples were dewaxed through three, five-minute interval pure xylene changes and rehydrated using a series of solutions of decreasing ethanol concentration; 100%, 90%, 70% for 3 minutes each followed by immersion in distilled water for 5 minutes. A hydrophobic pen was used to mark the area surrounding the tissue to minimise the solutions needed in the next steps. The tissue was then blocked using BSA for 1 hour at room temperature diluted in washing buffer. Using the same washing buffer, the tissue was washed several times and was incubated with the primary antibody (rabbit polyclonal to CD41 (abcam® ab63983), mouse monoclonal to CD41 [M148] (abcam® ab11024) or goat polyclonal to serotonin transporter (abcam® ab130130), rabbit polyclonal to collagen I (abcam® ab34710), goat polyclonal to alpha smooth muscle actin (abcam® ab21027)) for one hour at room temperature. Following taht, the samples were washed three times with PBS for 3-minute intervals and incubated with the secondary polyclonal antibodies (goat anti-rabbit alexa fluoro 488 (abcam® ab150077), donkey anti-mouse alexa fluoro 568 (abcam® ab175700) or a donkey anti-goat alexa fluor 555 (Thermofisher A-21432)) diluted in PBS to a concentration of 1:10 for 1 hour. The samples were then rewashed three times with PBS for 3-minute inter-vals and incubated with pure OsteoSense 680EX (PerkinElmer NEV10020EX) for 20 minutes. Two PBS washes of 10 minutes each were then carried out, and the tissue was incubated in DAPI stain (abcam® ab228549) at a concentration of 1:1000 (diluted in PBS) for 15 minutes. Each sample was then washed twice using PBS for 3-minute intervals. Finally, the samples were mounted with a coverslip using Fluoroshield mounting medium (abcam® ab104135) and sealed using nail polish at the edges of the coverslip.

For slight demineralisation of the whitlockite nanoparticles in the tissue, 0.5 M EDTA was applied to the samples for 30 minutes. The samples were then blocked for 1 hour using 5% BSA diluted in PBS. Samples were then washed three times with PBS for 3-minute intervals and incubated with one undiluted primary antibody (rabbit polyclonal to CD41 (abcam® ab63983), mouse monoclonal to CD41 [M148] (abcam® ab11024) or goat polyclonal to serotonin transporter (abcam® ab130130), rabbit polyclonal to collagen I (abcam® ab34710), goat polyclonal to alpha smooth muscle actin (abcam® ab21027)) for one hour at room temperature. Following that, samples were washed three times with PBS for 3-minute intervals and incubated with the secondary polyclonal antibodies (goat anti-rabbit alexa fluoro 488 (abcam® ab150077), donkey anti-mouse alexa fluoro 568 (abcam® ab175700) or a donkey anti-goat alexa fluor 555 (Thermofisher A-21432)) diluted in PBS to a concentration of 1:10 for 1 hour. The samples were then rewashed three times with PBS for 3-minute intervals and incubated with pure OsteoSense 680EX (PerkinElmer NEV10020EX) for 20 minutes. Two PBS washes, 10 minutes each were then carried out, and the tissue were incubated in DAPI stain (abcam® ab228549) at a concentration of 1:1000 (diluted in PBS) for 15 minutes. The samples were then washed twice using PBS for 3-minute intervals. Finally, samples were mounted with a coverslip using Fluoroshield mounting medium (abcam® ab104135) and sealed using nail polish at the edges of each coverslip.

For the osteogenic and neutrophil markers, aorta tissue samples were incubated as previously described with conjugated primary antibodies: Anti-CD11b antibody (abcam® ab204471), Anti-RUNX1/AML1+RUNX3+RUNX2 (abcam® ab207253), Anti-Osteopontin (abcam® ab282279), and IVISense Osteo 750 Fluorescent Probe.

All samples were imaged using an Olympus FV1000 and a Zeiss LSM 980 Airyscan confocal microscope. All antibodies were tested for specificity using secondary antibody and negative controls where appropriate.

Co-localisation analysis was done using the Coloc 2 function of FIJI Image J software, which was obtained as the Mandres' overlap coefficient (330, 331).

***Protein extraction from calcified particles:*** After isolation of the calcified particles, these

were incubated in 0.5 M EDTA for 24 hours at room temperature and constant agitation. They were then centrifuged at 20238 rcf for 10 minutes, the supernatant was collected, and the pellet was re-suspended in EDTA for 72 hours at room temperature and constant agitation. The final solution and the 24-hour supernatant were both used in SDS PAGE and dot blot. Following demineralisation, the resulting solution was added to a triton lysis buffer (15 0mM NaCl, 1% Triton X-100 and 50 mM Tris pH 8). It was added in one portion (sample-buffer ratio 1:1), which was incubated for 90 minutes at 4 ºC and then centrifuged at 16000 rcf for 10 minutes. The supernatant was collected, and the pellet discarded. To ensure the presence of proteins within the lysates, a Bradford assay was carried out using 50 μl of the sample (325). Following that, demineralised particles were added to a 2x laemmli sample buffer (65.8 mM Tris-HCl, pH 6.8, 26.3% (w/v) glycerol, 2.1% SDS (Bio-rad #1610737) with added 50 mM dithiothreitol (DTT)) (sample- buffer ratio 1:1). The solutions were then boiled at 95 ºC for 5 minutes and centrifuged at 16000g for 5 minutes. The supernatant was collected and either used immediately or aliquoted and stored at - 80 ºC for several months.

***Dot blot:*** Proteins extracted from platelet and particles were deposited onto a membrane. Five drops of 2 μl were used; each drop was left to dry prior to the addition of the next drop. For the serotonin and collagen controls, pure serotonin (Sigma Aldrich 14927) and collagen I (Sigma Aldrich C3867) samples were used. The membrane was then blocked using 5% dry milk powder for one hour at room temperature; the drops were changed every 15 minutes to ensure no precipitation of milk was taking place and that the whole membrane was blocked efficiently. The membranes were washed 4 times with PBS for 2 minutes each, incubated with primary antibodies (rabbit polyclonal to CD41 (abcam® ab63983), goat polyclonal to serotonin transporter (abcam® ab130130), rabbit monoclonal [EPR12735] to serotonin transporter (abcam® ab181034), rat monoclonal [YC5/45] to serotonin (abcam® ab6336), rabbit polyclonal to CD63 (abcam® ab118307), anti-CD62P antibody [EPR1444(2)(B)] (abcam®ab178424), rabbit polyclonal to collagen I (abcam® ab34710)) at a dilution of 1:10 in PBS for one hour. The membranes were then rewashed 4 times for 2 minutes each using PBS and were incubated with the secondary antibodies (goat anti-rabbit alexa fluoro 488 (abcam® ab150077), donkey anti-mouse alexa fluoro 568 (abcam® ab175700), anti-rat IgG (H+L) highly cross-ad-sorbed, CF™ 633 (Sigma Aldrich SAB4600133) or a donkey anti-goat alexa fluor 555 (Thermofisher A-21432)) at a concentration 1:100 in PBS for one hour. Finally, the membrane was rewashed 4 times for 2 minutes each and was imaged using a Leica optical microscope at a 10x magnification, 100 microsecond capture time, and 30% intensity.

***In vitro mineralization:*** Platelet-rich plasma was centrifuged at 6000g for 5 min and platelets were isolated. The platelet poor plasma was then centrifuged at 22000g for 15 min and serum was isolated. A solution of saturated at pH 5 was neutralized and Ph was adjusted to 7.4. Immediately the isolated platelets were mixed with collagen and gel was formed. The gel with platelets was them immersed in the serum and incubated at 37^o^C for up to 14 days. After 14 days, the gel was stained with Osteosense and Dapi, then imaged by SEM and fluorescence microscopy.

***Scanning electron microscopy (SEM) analyses and focused ion beam (FIB) sample preparation:*** Tissue samples or dewaxed histological sections were secured to an aluminium sample holder. Isolated calcified particles (and platelet dense granules) where suspended in deionised water and 5 µL was added to a clear glass slide secured to an aluminium sample holder with carbon tape. It was then silver painted and carbon coated in a Quorum K975X coater prior to imaging.

***SEM and energy-dispersive X-ray spectroscopy (EDX) analyses:*** Samples were SEM imaged using a Hitatchi S-3499N and a Carl Zeiss Crossbeam, operated at 10kV. Density-dependent colour SEM images were obtained as described in previous works[1]. Energy-dispersive X-ray spectroscopy (EDX) analysis was carried out using Oxford Instruments EDX detectors.

***Focused ion beam (FIB):*** A FEI Helios NanoLab 600 DualBeam Focused Ion Beam System was used. A micro region of each sample was coated with a Platinum layer at 30kV and 93pA. Following this procedure, currents between 93pA and 2.8nA were used for section lift out and thinning to 100nm.

***Transmission electron microscopy (TEM), scanning transmission electron microscopy (STEM), EDX and selected area electron diffraction (SAED) analyses:*** Samples extracted and prepared by FIB were imaged on a JEOL 2100 TEM. SAED were obtained at the TEM operating at 200 kV.

**Cell Isolation and Culture**

Human aortic smooth muscle cells were either purchased from ? or isolated from aortic tissue from donors at heart transplant, by explant. The collection of this tissue was approved by the local Ethics Committee. Briefly, tissue was collected in HBSS medium (Gibco L.T.I., Paisley, UK). After removal of the adventitia and surrounding connective tissue, the remaining intimal tissue was pinned onto tissue culture plastic (Sarstedt Ltd) using sterile 21-gauge needles, and was bathed in full VSMC medium: DMEM containing 1000 mg/l glucose (low glucose DMEM) supplemented with 2 mM l-glutamine, 100 u/ml penicillin/100μg/ml streptomycin, 25 mM HEPES pH 7.5, 1×non-essential amino acid solution and 15% FCS (Merck). Explant cultures were maintained at 37°C in a 5% CO2 atmosphere. VSMC were allowed to migrate out onto the plastic over a period of 2–8 weeks. Once there were large colonies of VSMC adhered to the plastic these were trypsinised and maintained as monolayers in Smooth muscle cell growth medium, (Cell Applications). Confluent monolayers of single isolates at between passages 3–10 were used for all experiments.

Human aortic endothelial cells were either purchased from Sigma-Aldrich or isolated from aortic tissue from donors at heart transplant, by enzymatic digestion. Tissue was collected in HBSS medium and subjected to collagenase H digestion (Roche) at 37oC for 15mins. Released endothelial cells were washed, centrifuged and resuspended in Endothelial Cell Growth Medium 2 (ECGM2 PromoCell) supplemented with 100 u/ml penicillin/100μg/ml streptomycin. Cells were plated into flasks coated with 1% bovine gelatin (Merck). Cells were passaged at confluency using trypsin/EDTA (Merck). Endothelial cell purity was verified by CD31 positivity by flow cytometry and isolates were used between passages 3-8.

**Cell Viability**

For cell experiments, these calcified particles were applied to a monoculture of HAECs and HASMCs by adding them directly into the media. Cell viability was determined by fluorescent labelling with a LIVE/DEAD Viability kit for mammalian cells (Invitrogen, US). 5000 cells were seeded on pre-coated chamber slides. After 24 hours, particles from different donors (healthy and diseased; with an O.D. of 1.0 and 50 µl per 500 µl of media) were added and further cultured for a week. After washing with PBS, cells were incubated with 2 µM calcein AM and 4 µM ethidium homodimer-1 for 30-45 minutes at RT and imaged using confocal microscopy (LSM 510 Meta inverted; Carl Zeiss).

**Cell Proliferation**

HAECs and HASMCs were seeded in a pre-coated 96 wells plate. Media was changed with 0.4% FCS in ECGM containing 150 U/ml P/S after attachment of the cells. After 24 hours, particles ((optical density of 1.0) 10 µl per 500 µl of media) were added and cultured for one week under normal conditions until further analysis. The proliferation assay was carried out with a CellTiter 96 AQueous Non-Radioactive Cell Proliferation Assay kit (Promega, US). Cells were incubated for one hour at 37 °C with 20 µl of MTS/PMS solution with 100 µl of media per well and absorbance was measured at 490 nm with a plate reader.

**Immunostaining**

HAECs and HASMCs, with and without particles, were cultured on coverslips for 3 weeks, washed two times with PBS, fixed in 4% formaldehyde solution (Sigma) for 10 minutes and washed three times with PBS to remove the fixative solution. The coverslips were permeabilized with Triton-X-100 (0.5% v/v, Sigma Aldrich) for 3 minutes and blocked for 30 min with BSA (3% w/v, Sigma). They were and then incubated with primary antibodies at room temperature for one hour, CD31 (Dako), CD144 (Dako), vWF (Dako), α-SMA (Dako), osteopontin (R&D) and Runx2 (R&D). Coverslips were washed and then incubated with VECTASTAIN® Elite® ABC-Peroxidase Kit (R.T.U. Universal, Vector Laboratories, 2BScientific, Kidlington, UK) for 30 min for each secondary and tertiary antibody (Vector Laboratories). Specimens were then incubated with DAB (Sigma) for 5 min, washed well in tap water, nuclei were stained with haematoxylin for 1 min, and slides were mounted using Aquatex (VWR). Digital images were taken on a Nanozoomer (Hamamatsu, Shizuoka, Japan).

**Immunohistochemistry**

Samples were washed with PBS, fixed in 10% formalin saline (Sigma) for 5-10 minutes and washed with deionized water. To assess calcium deposits, an Alizarin Red staining was used. Coverslips were stained in 2% Alizarin Red S solution for 5 minutes and mounted with DPX (VWR). Von Kossa stainings were used to visualize mineralization. Coverslips were incubated in 2% silver nitrate (VWR) solution for 1 hour under light conditions. Samples were then washed with deionized water, incubated in 2.5% sodium thiosulphate (VWR) for 5 minutes, washed in tap water and counterstained with van Giesson (VWR). Coverslips were mounted in DPX. Coverslips were analysed by brightfield microscopy with the Axio Scan.Z1 (Zeiss).

**Statistical Analysis**

Statistical analysis was performed using GraphPad Prism 5 (GraphPad Software, United States). Prior to analysis of significant differences, data was subjected to a Shapiro-Wilk normality test. Normally distributed data was subjected to a t-test or one-way ANOVA, followed by a Tukey post-hoc test. Other data was analysed by a Kruskall Wallis nonparametric test. Differences were considered significant for p < 0.05. Data is presented as mean ± standard deviation. For the Westerns, data is presented as median ± standard deviation.

**References**

1.	Bertazzo, S., et al., *Nano-analytical electron microscopy reveals fundamental insights into human cardiovascular tissue calcification.* Nat Mater, 2013. **12**(6): p. 576-583.
